# Microbiome and Antimicrobial Resistance Gene Dynamics in International Travelers

**DOI:** 10.1101/506394

**Authors:** Charles Langelier, Michael Graves, Katrina Kalantar, Saharai Caldera, Robert Durrant, Mark Fisher, Richard Backman, Windy Tanner, Joseph L. DeRisi, Daniel T. Leung

## Abstract

We engaged metagenomic next generation sequencing to longitudinally assess the gut microbiota and antimicrobial resistomes of international travelers to understand global exchange of resistant organisms. Travel resulted in an increase in antimicrobial resistance genes and a greater proportion of *Escherichia* species within gut microbial communities without impacting diversity.

## Background

International travel is a known contributor to the global emergence of antimicrobial resistant (AMR) organisms (1–4). Colonization with resistant microbes acquired during travel may persist asymptomatically for extended periods and result in transmission into the environment and susceptible populations (5). The mechanisms underlying acquisition of AMR bacteria during travel are incompletely understood, although changes in the intestinal microbiota are hypothesized to play a role (6). Here, we engaged metagenomic next generation sequencing (mNGS) to assess both gut microbiota composition and the antimicrobial resistome with a goal of understanding AMR bacteria acquisition among healthcare workers traveling internationally.

## Methods

Adults with planned travel to Asia or Africa for healthcare-related work were recruited between March 2016 and February 2018. Participants introduced one tablespoon of stool into vials with either RNAprotect (Qiagen) or Cary-Blair (CB) media and then submitted samples and surveys pre-travel (PRE), post-travel (PST), 30 days post-travel (30D), and 6 months post-travel (6MO). Upon receipt, RNAprotect samples were stored at −80°C, and CB samples at 4°C. CB samples were inoculated onto differential chromogenic agar (CHROMagar) plates and incubated overnight at 37°C. Single colonies were then inoculated into LB broth, and incubated overnight at 37°C. If multiple morphotypes were identified, separate subcultures were done. Speciation was performed using MALDI-TOF mass spectrometry.

DNA and RNA extracted using the Qiagen Powerfecal kit underwent metagenomic sequencing as previously described (7). Raw data are available publicly via Sequence Read Archive (SRA) accession number: SUB4474900. Detection of enteric microbiota leveraged a previously described bioinformatics pipeline (7). Microbial alignments were aggregated at the genus-level prior to downstream analyses. To control for background environmental and reagent contaminants, we sequenced no-template water control samples and directly subtracted the number of reads aligning to each genus present in these controls from other samples.

The SRST2 computational tool and Argannot2 database were used to identify AMR genes with allele coverage of at least 20%(8). While a precise definition of ESBL has not been established, we employed a working definition of Ambler class A-D β-lactamases with known or predicted ability to confer resistance to penicillins and first through third-generation cephalosporins and resistant to clavulanic acid (9,10). We required detection of chromosomally-encoded class C beta lactamases (ie. *AmpC*) by both DNA-Seq and RNA-Seq to capture actively expressed genes.

## Results

Nine of 10 subjects were culture-positive for ESBL-producing *Escherichia coli* upon return, including eight who traveled to Nepal and one who went to Nigeria. One subject was found to be colonized prior to departure (T3), three had persistent ESBL-PE carriage at the 30-day visit (T2, T3, T5) and two at six months (T3, T5) (**Table 1**). While four subjects experienced diarrheal symptoms during travel, only one (T5) had persistent diarrheal symptoms at six months. Diarrheal symptoms were not associated with persistent ESBL-PE colonization at any time point and no travelers reported antibiotic use.

**Table 1.**
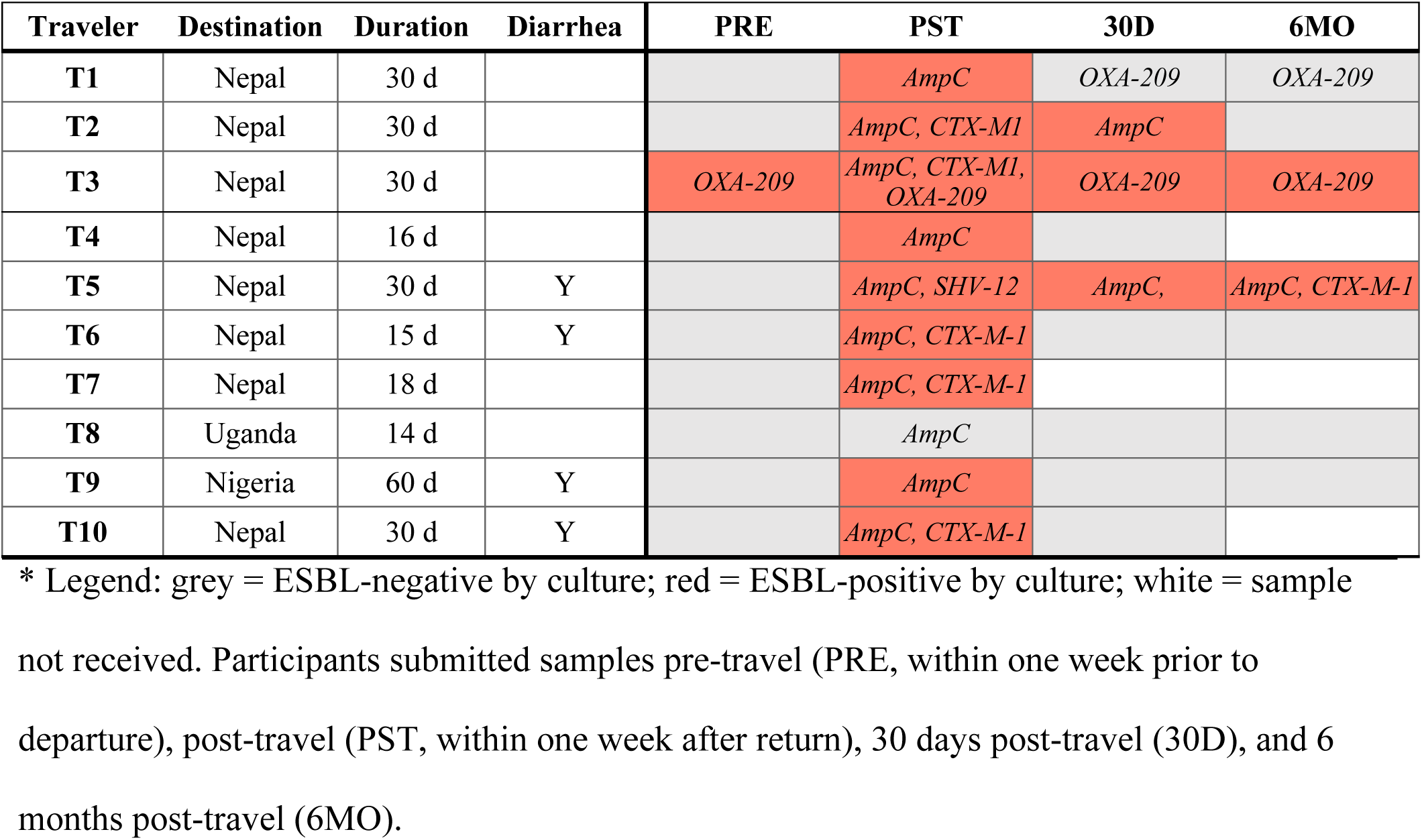
Traveler characteristics and assessment of extended spectrum beta lactamases.

| Traveler | Destination | Duration | Diarrhea | PRE | PST | 30D | 6MO |
| --- | --- | --- | --- | --- | --- | --- | --- |
| T1 | Nepal | 30 d |  |  | *AmpC* | *OXA-209* | *OXA-209* |
| T2 | Nepal | 30 d |  |  | *AmpC, CTX-M1* | *AmpC* |  |
| T3 | Nepal | 30 d |  | *OXA-209* | *AmpC, CTX-M1, OXA-209* | *OXA-209* | *OXA-209* |
| T4 | Nepal | 16 d |  |  | *AmpC* |  |  |
| T5 | Nepal | 30 d | Y |  | *AmpC, SHV-12* | *AmpC,* | *AmpC, CTX-M-1* |
| T6 | Nepal | 15 d | Y |  | *AmpC, CTX-M-1* |  |  |
| T7 | Nepal | 18 d |  |  | *AmpC, CTX-M-1* |  |  |
| T8 | Uganda | 14 d |  |  | *AmpC* |  |  |
| T9 | Nigeria | 60 d | Y |  | *AmpC* |  |  |
| T10 | Nepal | 30 d | Y |  | *AmpC, CTX-M-1* |  |  |

\* Legend: grey = ESBL-negative by culture; red = ESBL-positive by culture; white = sample not received. Participants submitted samples pre-travel (PRE, within one week prior to departure), post-travel (PST, within one week after return), 30 days post-travel (30D), and 6 months post-travel (6MO).

We first examined changes in gut microbiome alpha diversity following international travel (**Figure 1A**) and found that SDI did not significantly change upon return or at day 30 post-travel (P = 0.674 and 0.250, respectively, by Wilcoxon Rank Sum. We next asked whether microbial community composition differed across all subjects at their post-travel visit versus pre-travel but found no difference (Bray Curtis Index, P = 0.23 by PERMANOVA). Even though global composition and diversity of gut microbiota didn’t significantly change following travel, significant differences in the abundance of discrete genera were observed (**Figures 1B** and **Appendix Figure 1**). Across all subjects, *Enterobacteriaceae* demonstrated the greatest fold change in abundance post-travel, and of these *Escherichia* was the genus most differentially increased (P < 0.001 by Wilcoxon Rank Sum) (**Figures 1B and Appendix Figure 1**).

**Figure 1.**
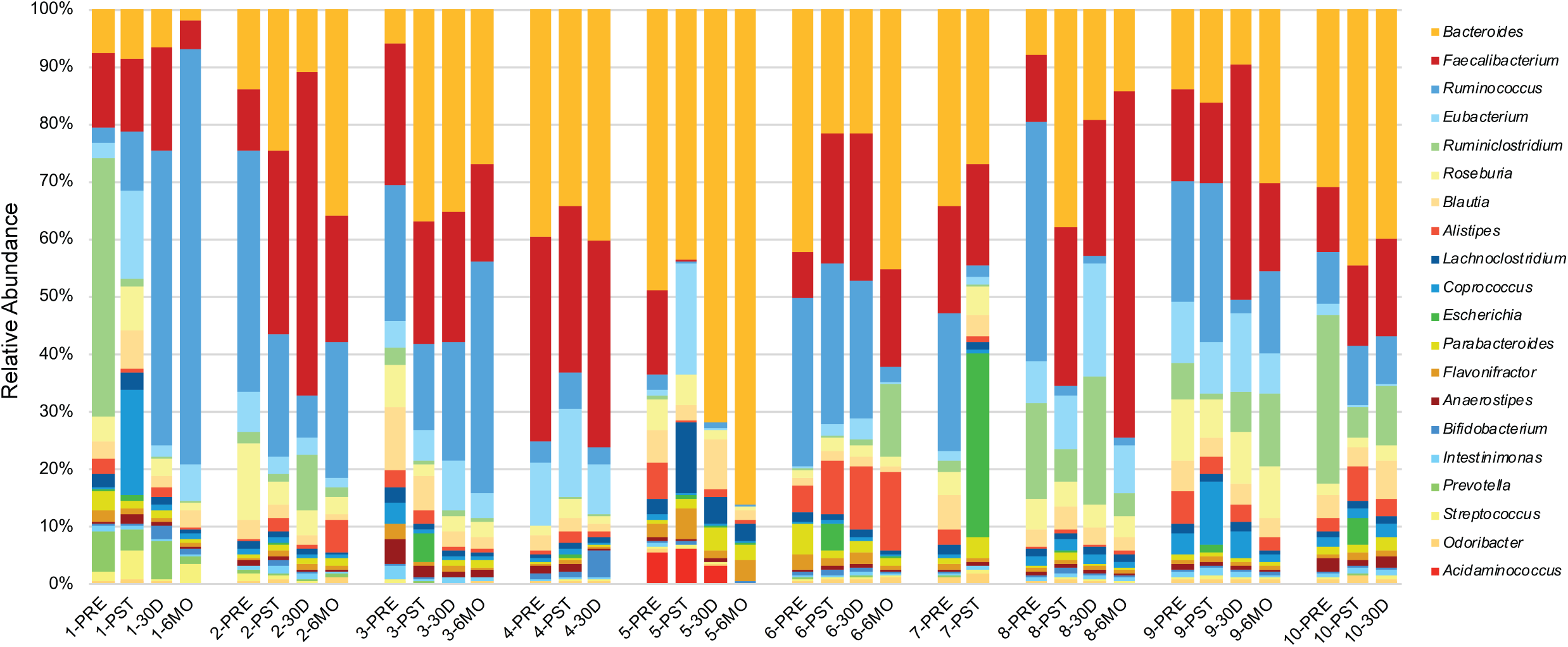
A) Longitudinal profile of traveler gut microbiome diversity measured by Shannon Diversity Index (SDI). Subject T5 had an SDI more than three standard deviations below the mean when measured at 30 days and 6 months. B) Microbes, by genus, demonstrating the greatest fold change in abundance following travel based on DNA-Seq nucleotide alignments. C) Total number of AMR genes identified with at least 20% allele coverage by DNA-Seq. D) Total number of AMR genes identified with at least 20% allele coverage by RNA-Seq. <u>Legend</u>: BLA: beta-lactamase, SUL: sulfa, GLY: glycopeptide, FLQ: fluoroquinolone, DFR: dihydrofolate reductase, AGL: aminoglycoside, MLS: macrolide, lincosamide, streptogramin, Tet: tetracycline, ESBL: extended-spectrum beta-lactamase.

Interrogation of the antimicrobial resistome revealed an increase in AMR genes and transcripts following return from travel (P < 0.01 for DNA-Seq and P = 0.03 for RNA-Seq, respectively, by Wilcoxon rank sum, **Table 2, Figures 1C-D, Appendix Figure 2**). ESBL and/or *AmpC* encoding genes were identified in 100% of samples with an ESBL culture-determined phenotype and 14% of samples without, including the single subject (T8) who was phenotypically ESBL-negative following travel (**Table 1**). Beta-lactam resistance genes increased post-travel including *AmpC, CTX-M, OXA and SHV* gene families known or predicted to confer ESBL production, as well as diverse additional beta lactamase genes (**Appendix Figure 2**). Travel also resulted in an increase in *qnr* plasmid-mediated quinolone resistance genes, as well as trimethoprim (*dfr*), sulfa, macrolide and aminoglycoside resistance genes (**Table 2**). Genes conferring resistance to tetracyclines and aminoglycosides were most abundant in subjects at baseline and remained stable or decreased during travel (**Table 2**).

**Table 2.** Fold change in abundance of AMR genes detected by DNA-Seq with at least 20% allele coverage at the pre-travel and post-travel visits, listed by drug class or antimicrobial resistance gene class.

| Antibiotic<br>class or<br>AMR gene | Fold Change<br>Pre- vs. Post-Travel |  |
| --- | --- | --- |
|  | DNA-Seq | RNA-Seq |
| BLA | 6 | 9 |
| -*AmpC* | >100 | >100 |
| -*AmpH* | >100 | >100 |
| -*CTX* | >100 | >100 |
| -*MrdA* | >100 | >100 |
| -*OXA* | 2 | 1 |
| -*SHV* | >100 | >100 |
| -*TEM* | >100 | >100 |
| FLQ | >100 | >100 |
| TMP | >100 | >100 |
| SMX | 21 | 29 |
| MLS | 2 | 7 |
| TET | <1 | 1 |
| GLY | <1 | <1 |
| AGL | 2 | 5 |

\* Legend: BLA: beta-lactamase, SUL: sulfa, GLY: glycopeptide, FLQ: fluoroquinolone, DFR: dihydrofolate reductase, AGL: aminoglycoside, MLS: macrolide, lincosamide, streptogramin, Tet: tetracycline, ESBL: extended-spectrum beta-lactamase.

| Antibiotic class or AMR gene | Fold Change<br>Pre- vs. Post-Travel |  |
| --- | --- | --- |
|  | DNA-Seq | RNA-Seq |
| BLA | 6 | 9 |
| -AmpC | >100 | >100 |
| -AmpH | >100 | >100 |
| -CTX | >100 | >100 |
| -MrdA | >100 | >100 |
| -OXA | 2 | 1 |
| -SHV | >100 | >100 |
| -TEM | >100 | >100 |
| FLQ | >100 | >100 |
| TMP | >100 | >100 |
| SMX | 21 | 29 |
| MLS | 2 | 7 |
| TET | <1 | 1 |
| GLY | <1 | <1 |
| AGL | 2 | 5 |

**Figure 2.**
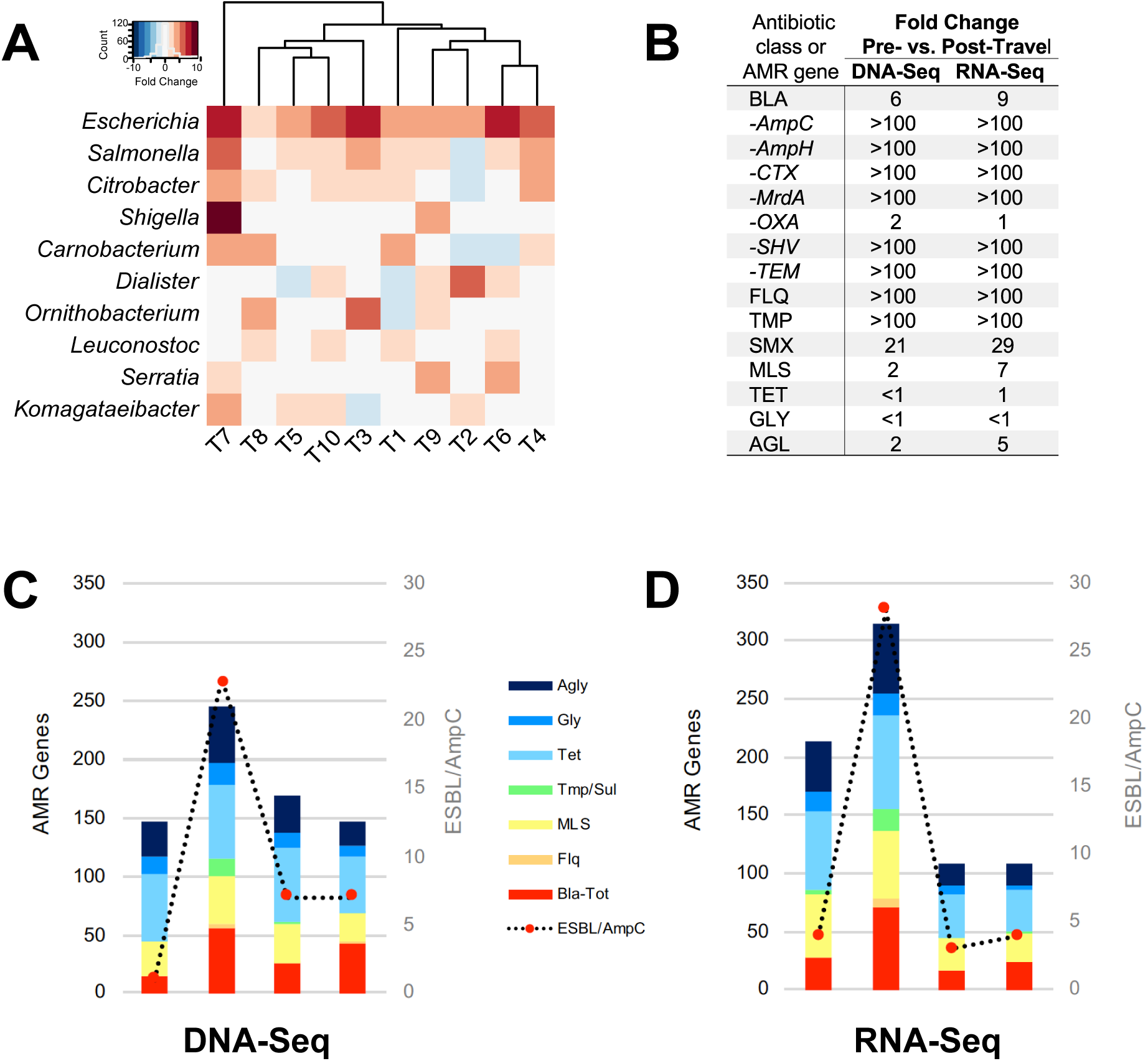
Relative proportion of the 20 most abundant microbial genera present in the enteric microbiomes of international travelers, assessed longitudinally using shotgun metagenomic DNA sequencing. Participants submitted samples pre-travel (PRE, within one week prior to departure), post-travel (PST, within one week after return), 30 days post-travel (30D), and 6 months post-travel (6MO).

We found no significant differences in SDI between persistent carriers and those who lost carriage 30 days (n=3, P = 0.56 by t-test) or at six months (n=2, P = 0.27 by t-test). T5, who was colonized at both time points and who was the only subject with persistent diarrheal symptoms, had an SDI more than three standard deviations below the mean at six months (**Figure 1A**). Bray-Curtis distance measured pre- or post-travel did not differ between subjects who were ESBL-PE positive by culture at 30 days or six months (P = 0.32 by PERMANOVA). No taxa were associated with post-travel ESBL-positivity based on an adjusted P value < 0.05 at 30 days or six months.

## Discussion

International travel is a well-recognized contributor to the global spread of emerging infectious diseases, including AMR bacteria (1,4). Here, we interrogated the enteric microbiota and resistomes of returned travelers and found a marked increase in AMR genes that was associated with an increased proportion of *Escherichia*. ESBL and actively transcribed *AmpC* genes were amongst the most increased AMR genes post-travel, in line with prior reports (4). Notably both mNGS and culture-based methods demonstrated persistent ESBL colonization after six months in 20% of subjects, confirming prior work demonstrating that travel can induce long-term changes in the antimicrobial resistome (5). In addition to beta-lactamase genes, mNGS identified a diversity of other AMR gene classes that increased in abundance following travel. For instance, 80% of subjects acquired horizontally-transferrable *qnr* genes following travel, reflecting the limited utility of quinolones for treatment of traveler’s diarrhea (11).

Changes in microbiome diversity were not associated with ESBL-positivity at 30 days nor at six months, suggesting that disruption of the antimicrobial resistome can occur in the setting of a preserved microbial community structure. We observed a high rate of ESBL-PE acquisition in this cohort, the majority of whom traveled to the Indian subcontinent. This observation is consistent with published reports of ESBL-PE acquisition rates of up to 90% in travelers returning from this region, in particular those who took antibiotics (1). Notably, none of the travelers in this cohort reported antibiotic use, suggesting that significant ESBL-PE colonization can occur even in the absence of antibiotic-related disruption of commensal gut microbiota.

This study is limited by small sample size, and as such, relevant associations may have been missed. Future studies leveraging mNGS to assess larger cohorts traveling to a greater diversity of destinations are needed. Nonetheless, these results highlight the pervasiveness of AMR microbe exchange during international travel and the promise of mNGS for assessing the global exchange of antimicrobial resistance.

## Author Bio

(first author only, unless there are only 2 authors)

Dr. Langelier is an Assistant Professor in the Division of Infectious Diseases at the University of California, San Francisco. His research interests involve using metagenomics and transcriptional profiling to investigate host–pathogen interactions and understand the causes of diagnostically challenging diseases.

**Appendix Figure 1.** Assessment of relative abundance of *Escherichia* over time, pre- and post-travel, revealed a statistically significant increase (P < 0.001 by Wilcoxon Rank Sum) post-travel.

**Appendix Figure 2.** Beta-lactamase genes identified in the gut microbiomes of travelers over time, pre- and post-travel. Gradient shading corresponds to gene abundance (average allele sequencing depth per million reads mapped, AdpM).

