## Supplementary figures and images for "Microbiome and Antimicrobial Resistance Gene Dynamics in International Travelers"

### Supplemental Table 1

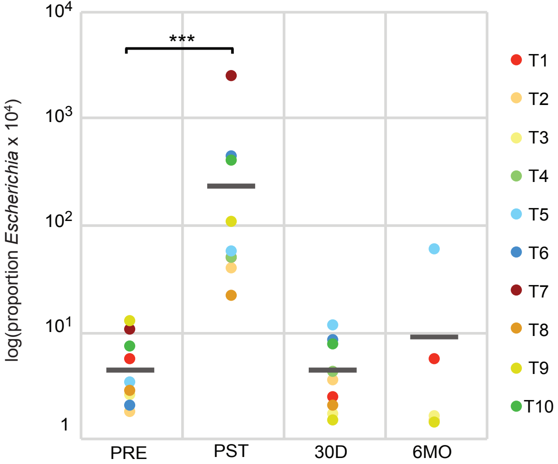

### Supplemental Table 2

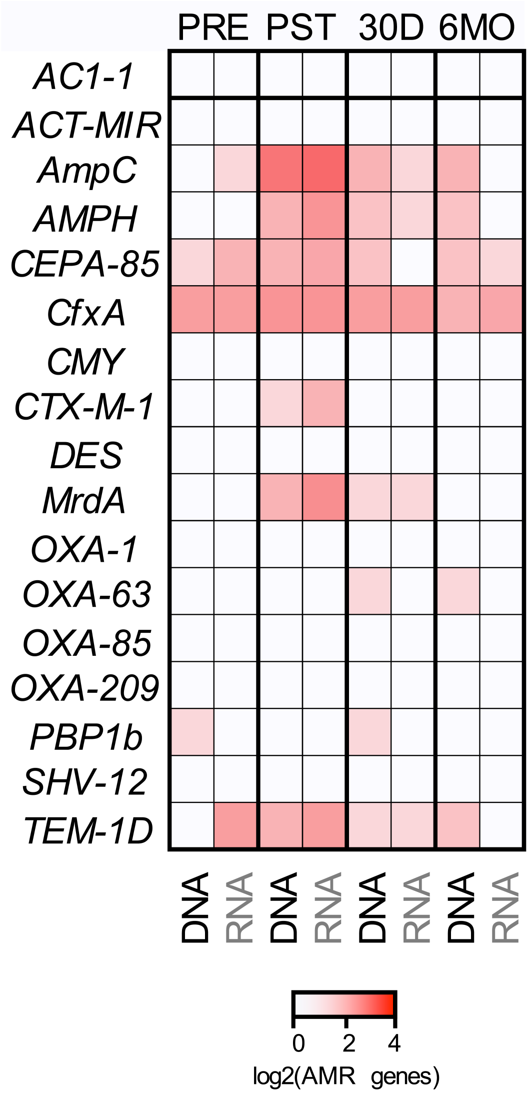
